# β-ionone regulates *Arabidopsis thaliana* transcriptome and increases its resistance against *Botrytis cinerea*

**DOI:** 10.1101/2023.05.02.539130

**Authors:** Abrar Felemban, Juan C. Moreno, Jianing Mi, Shawkat Ali, Arjun Sham, Synan F. AbuQamar, Salim Al-Babili

**Author notes:** Corresponding authors: Synan F. AbuQamar and Salim Al-Babili. Equal contribution (Abrar Felemban, Juan C. Moreno).

## Abstract

Carotenoids are isoprenoid pigments vital for photosynthesis. Moreover, they are the precursor of apocarotenoids that include the phytohormones abscisic acid (ABA) and strigolactones (SLs), and retrograde signaling molecules and growth regulators, such as β-cyclocitral and zaxinone. The apocarotenoid β-ionone (β-I) was previously reported to exert antimicrobial effects. Here, we showed that the application of this scent to Arabidopsis plants at micromolar concentrations caused a global reprogramming of gene expression, affecting thousands of transcripts involved in stress tolerance, growth, hormone metabolism, pathogen defense and photosynthesis. These changes, along with modulating the levels of the phytohormones ABA, jasmonic acid and salicylic acid, led to enhanced Arabidopsis resistance to *Botrytis cinerea* (*B.c.*), one of the most aggressive and widespread pathogenic fungi affecting numerous plant hosts and causing severe losses of postharvest fruits. Pre-treatment of tobacco and tomato plants with β-I followed by inoculation with *B.c.* confirms the conserved effect of β-I and induced immune responses in leaves and fruits. Moreover, there was reduced susceptibility to *B.c.* in *LYCOPENE β-CYCLASE-* expressing tomato fruits possessing elevated levels of the endogenous β-I, indicating beneficial biological activities of this compound *in planta*. Our work unraveled β-I as a further carotenoid-derived regulatory metabolite and opens up new possibilities to control *B.c.* infection by establishing this natural volatile as an environmentally friendly bio-fungicide.

## Introduction

Carotenoids are a large group of isoprenoid pigments that include more than 1000 distinct compounds (Yabuzaki, 2017). The conjugated double bond of carotenoids, which is responsible for their color and functions in photosynthesis, makes them prone to oxidative cleavage caused by attacks of reactive oxygen species (ROS) or through enzymatic reactions catalyzed by CAROTENOID CLEAVAGE DIOXYGENASES (CCDs) (Giuliano et al., 2003; Beltran and Stange, 2016; Hou et al., 2016; Moreno et al., 2021). This metabolic process gives rise to a family of important metabolites called apocarotenoids, which includes the precursors of abscisic acid (ABA) and strigolactones (SLs), two plant hormones with diverse biological functions ranging from rhizospheric communications and regulating seed dormancy to biotic and abiotic stress response (Al-Babili and Bouwmeester, 2015; Moreno et al., 2021). Moreover, some apocarotenoids, such as β-cyclocitral (β-cc), β-cyclocitric acid, zaxinone, anchorene and iso-anchorene, are retrograde signals in the plastid-nucleus communication, acting as growth regulators and mediating plant response to oxidative stress (Ramel et al., 2012; D’Alessandro et al., 2018; D’Alessandro et al., 2019; Dickinson et al., 2019; Jia et al., 2019; Wang et al., 2019; Jia et al., 2021).

β-ionone (β-I) is a C_13_ volatile compound formed from β-carotene by ROS (Fig. S1A) during photosynthesis, particularly under high light conditions (Ramel et al., 2012). In addition, it is produced by several carotenoid cleavage dioxygenases (CCDs) that cleave the C9-C10 and/or C9′-C10′ double bond(s) in β-carotene (Fig. S1B-D) (Auldridge et al., 2006; Alder et al., 2012; Bruno et al., 2015). CCD1 targets these two double bonds, producing two C_13_ volatile compounds β-I and a C_14_ dialdehyde (Fig. S1B) (Wei et al., 2011), while Arabidopsis CCD4 mediates a single cleavage reaction in β-carotene, leading to all-*trans*-β-apo-10’-carotenal (C_27_) and one molecule of β-I (Fig. S1C) (Rubio-Moraga et al., 2014; Bruno et al., 2015; Bruno et al., 2016). CCD7 is a stereospecific enzyme that cleaves 9-*cis*-β-carotene into 9-*cis*-β-apo-10′-carotenal (C_27_) and β-I (Fig. S1D) (Alder et al., 2012; Bruno et al., 2014; Bruno et al., 2016; Haider et al., 2018). β-I has been implicated in biotic stress response against herbivores (Griffin et al., 1999; Wei et al., 2011). Moreover, it was reported to be an insect repellant and to have antibacterial and fungicidal properties (Giuliano et al., 2003; Caceres et al., 2016; Aloum et al., 2020), inhibiting growth of *Peronospora tabacina* (Salt et al., 1986), *Candida albicans* (Griffin et al., 1999), *Aspergillus niger* (Hassan and Bakhiet, 2017), and *Colletotrichum musae* (Utama et al., 2002).

*Botrytis cinerea* (*B.c.*), the causal agent of gray mold disease, is considered one of the most destructive fungal pathogens due to its capability of infecting over 200 different plant species (Govrin and Levine, 2000; Shlezinger et al., 2011; Valeri et al., 2021). Despite the employment of chemical management for many years, the capacity of *B.c.* to quickly adapt to chemical pesticides has made it a recurrent issue (Rosslenbroich and Stuebler, 2000; Zhao et al., 2010; Panebianco et al., 2015). This phenomenon and the ecological impact of chemical fungicides have proposed novel alternatives of disease management, including bio-fungicides. Interestingly, several natural compounds can activate plant defense response, by inducing a physiological condition that activates defense response upon subsequent stress (Conrath, 2009; Aranega-Bou et al., 2014; Li et al., 2022). Therefore, finding natural compounds effective on plants to better cope with abiotic or biotic stresses appears a promising, sustainable strategy for disease control and/or alleviating the damage caused by abiotic stress.

In this study, we set out to explore if β-I acts as a regulatory metabolite, similarly to β-cc that arises from the same precursor and under similar conditions (Ramel et al., 2012), and to investigate whether it is involved in plant defense against the necrotrophic pathogen, *B.c.* By combining phenotypic, transcriptomic, and metabolomic analyses, we showed that β-I is a regulatory metabolite in Arabidopsis, which enhances the resistance to *B.c.* by provoking a transcriptional response overlapping with that triggered upon *B.c.* infection, and modulating different defense pathways, including those of ABA, salicylic acid (SA) ethylene (ET), and jasmonic acid (JA). The inhibitory effect of β-I against *B.c.* on leaf and fruit tissues was found to be conserved in dicotyledonous crops *i.e.,* tobacco and tomato. Moreover, transgenic tomato fruits with enhanced β-I content showed increased resistance to *B.c*, suggesting a potential application of β-I in crop production with a focus on reducing disease and pest incidence for achieving agricultural sustainability and food security.

## Results

### Exogenous β-ionone application modulated the expression level of defense- and growth-related genes

To get insights into the biological functions of β-I, we employed an RNAseq approach to explore possible transcriptional changes upon its application. For this purpose, we sprayed Arabidopsis plants twice with β-I and collected RNA samples at 3 and 24 hours post treatment (hpt; Fig. S2A), covering short and middle term transcriptional changes. Subsequent RNAseq analysis revealed a significant change in the transcript level of a striking number of genes (Dataset S1-S2), which was more pronounced at 24 hpt. Venn diagrams showed the upregulation of 1239 differentially expressed genes (DEGs) in response to β-I at early and late time points (common to both time points), and 684 and 5148 DEGs that were upregulated only at 3 and 24 hpt, respectively (Fig. 1A; see extended and full list in Figs. S4 and S5 and Dataset S3 and S5). We also detected a total of 907 downregulated DEGs at early and late time points, and 485 and 5301 DEGs that were downregulated only at 3 and 24 hpt, respectively (Fig. 1A; see extended and full list in Figs. S4 and S6 and Dataset S4 and S6). To understand the primary functions of these DEGs, we performed a gene ontology (GO) enrichment analysis (https://www.arabidopsis.org/tools/go_term_enrichment) to determine biological processes enriched in our dataset (Fisher’s exact test with FDR correction *p*<0.05). Our analysis revealed a high number of biological processes associated with the 1239 upregulated DEGs (Fig. S5; Dataset S3 and S5), including defense and response to fungal pathogens and to ABA/JA/ET/SA (Fig. 1A and Figs. S4 and S5). Part of the downregulated genes, at both time points, were involved in growth, development, and photosynthetic processes (Figs. S4 and S6; Dataset S4 and S6). Next, we used SUBA4 (Hooper et al., 2014; Hooper et al., 2017) to predict the localization of the proteins encoded by DEGs upon β-I treatment. Compared to the protein localization in non-treated Arabidopsis, the relative compartment distribution of proteins encoded by β-I up-regulated genes was high in the nucleus and cytosol, and lower in chloroplasts, with ratios of 32%, 21% and 8%, compared to 28%, 18% and 14%, respectively (Fig. 1B; Dataset S7-S9). The subcellular distribution of proteins encoded by the downregulated genes showed opposite patterns *i.e.,* they were less present in the nucleus (from 28% to 24%) and cytosol (from 18% to 16%), but more frequent in chloroplasts (from 14% to 21%) and mitochondria (from 9% to 12%), than that in the normal distribution. Then, we performed a MapMan analysis (Thimm et al., 2004) to unveil major changes induced by β-I. We observed that 34 out of 35 MapMan Bins were affected by β-I treatment (Dataset S10). The majority of these changes occurred in processes such as photosynthesis (128 genes), cell wall biosynthesis (185 genes), secondary metabolism (176 genes), hormone metabolism (253 genes), biotic stress (407 genes), RNA metabolism (1393 genes), DNA metabolism (322 genes), and signaling (598 genes) (Fig. 1C; Figs. S7-S8 and Dataset S10). We observed a clear pattern of repressing processes contributing to plant growth, such as photosynthesis, tetrapyrrole, chlorophyll, and isoprenoid biosynthesis (Fig. S9A-B). For instance, β-I decreased the expression of 97 out of 128 genes involved in photosynthesis, including light harvesting complexes (LHCs) genes, such as *LHCAs* and *LHCBs* (1.63 to 67-fold), *RuBisCO small subunit 1A*/*RBCS1A* and *RBCS1-3B* (2.7 to 46-fold). Moreover, it repressed genes of photosystem subunits, such as *PSAG* (5.1-fold), *PSB28* (20-fold), *PSBQ* (73-fold), *PSBS* (27-fold), and subunits of the plastoquinone dehydrogenase and ATPase complexes, such as *NDH-M* (57-fold) and *ATPD* (6.9-fold) (Fig. 1C; Fig. S9B). Out of 35 genes, the transcript of 29 genes involved in tetrapyrrole and chlorophyll biosynthesis, including *glutamyl-tRNA reductase/HEMA3* (30-fold), *protochlorophyllide oxidoreductase*/*PORC* (18.3-fold), and *chlorophyll synthetase* (11.7-fold), *genomes uncoupled4*/*GUN4* (2.94-fold) and *GUN5* (4.6-fold) were also reduced when plants were treated with β-I (Dataset S10). The majority of genes involved in the methylerythritol phosphate (MEP) pathway, which provides the precursor for chlorophylls and carotenoids, were also downregulated (2.1-5.4-fold; Fig. S9A and Dataset S10). By contrast, genes related to plant defense comprising processes, such as hormone metabolism, biotic stress response, RNA metabolism, and signaling showed high increases upon β-I treatment (Fig. 1C). For instance, we observed an induction of *PROTEASE INHIBITOR* (*PI*; *at4g12470*; 13.4-fold), *PLANT DEFENSIN1.4* (*PDF1.4*; 14.1-fold), *NIMIN-1-related* (2.5-fold), *RECEPTOR LIKE PROTEIN* (*RLP22*; 854-fold), and *DEFENSIN-LIKE PROTEINs* (*DEFLs*; 5 to 204-fold). Moreover, representatives of several transcription factor families, such as *WRKYs* (2.1-1585-fold), *ANACs* (34.5-43.7-fold), *ERFs* (1.5-27-fold), and *RAPs* (1.4-82.2-fold) were upregulated. We observed enhanced gene expression for genes involved in signaling such as *MPKs* (1.8-2.8-fold), *ENHANCED DISEASE RESISTANCE1* (*EDR1*; 2-fold), and RESPONSIVE TO *DESICCATION 20* (*RD20*; 18.44-fold).

**Fig. 1.**
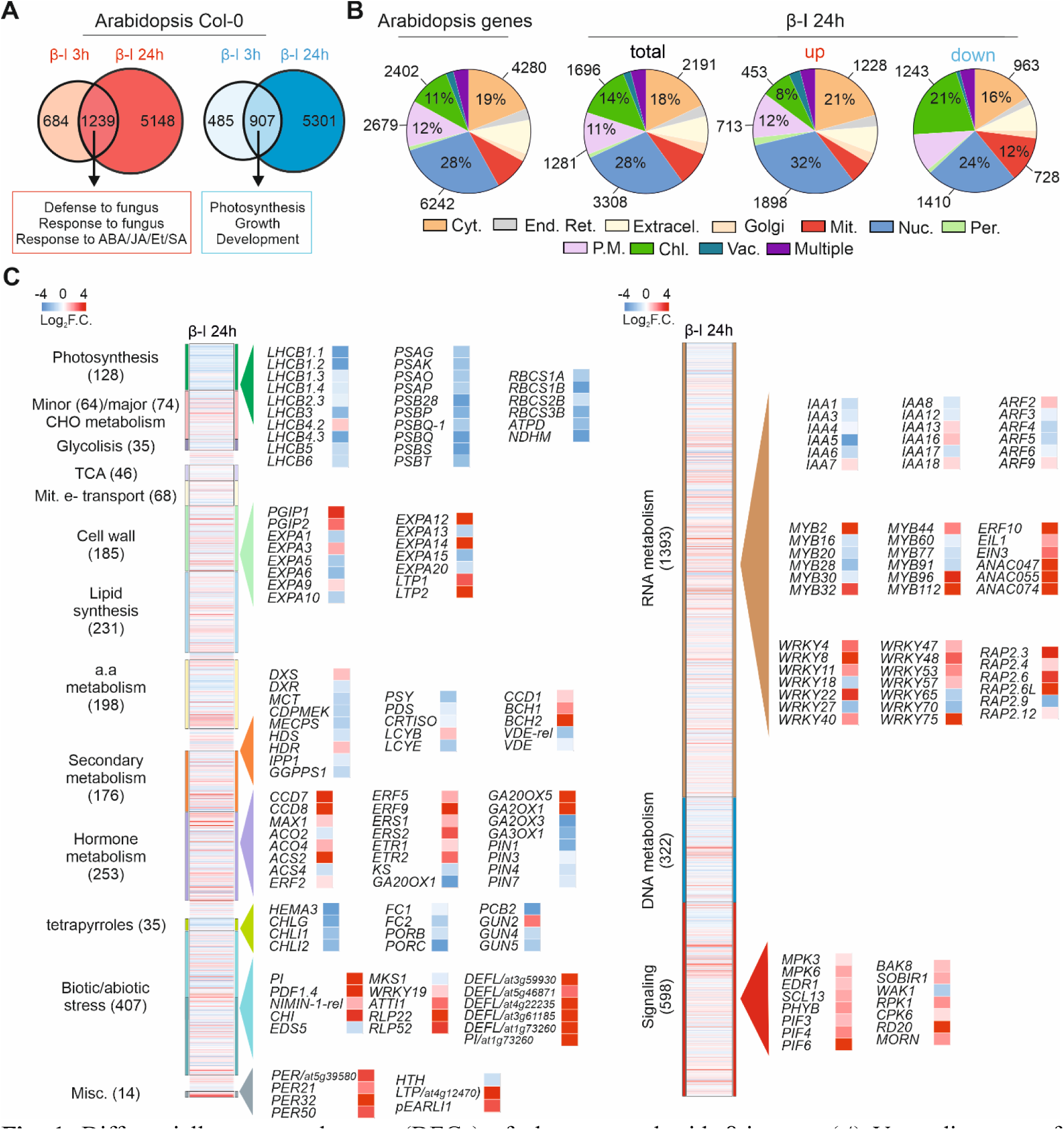
Differentially expressed genes (DEGs) of plants treated with β-ionone. (*A*) Venn diagrams of DEGs identified through RNAseq in Arabidopsis plants treated with β-I at 3 and 24 hpt. Venn diagrams representing up- and downregulated genes are shown in red and blue, respectively. Some of the most important GO biological processes enriched in the overlapping genes are shown (Table S1; Fig. S4). (*B*) Distribution of cellular compartments for genes present in the reference Arabidopsis genome (TAIR) and DEGs in the β-I treatment at 3 and 24 hours post treatment (hpt). Subcellular localization analysis was performed in SUBA4 online software (https://suba.live/) (Dataset S7-S9 for full subcellular localization list). Genes encoding proteins with 2 or more localizations were grouped in the “multiple” category. Each color in the pie chart represents a cellular compartment. (*C*) Heatmap representation of transcriptional changes in Arabidopsis plants treated with β-I at 24 hpt. Heatmaps show 15 MapMan bins with profound transcriptional changes (Dataset S10 for full list and description of each gene). Statistical analysis for the RNAseq was performed using DESeq2 with Benjamini and Hochberg’s approach for controlling False Discovery Rate (FDR). Genes were adjusted Log_2_ fold change expression (padj<0.05). Cyt: cytoplasm; End. Ret.: Endoplasmic reticulum; Extracel.: extracellular; Mit.: mitochondria; Nuc.: nucleus; Per.: peroxisome; P.M.: plasma membrane; Chl.: chloroplast; Vac.: vacuole.

### β-ionone enhanced Arabidopsis resistance against the necrotrophic fungus *Botrytis cinerea*

In our RNAseq analysis, we noticed that β-I induced the expression of genes involved in plant defense and repressed those required for growth, indicating the possibility that this apocarotenoid may contribute to the plant defense against pathogens. This hypothesis is supported by previous studies reporting on anti-microbial and anti-fungal properties of β-I; although at very high concentrations of millimolar ranges (Ozaki et al., 2008; Harada et al., 2009). Taking into consideration the importance of the necrotrophic fungus *B.c.* for basic science and agriculture, we tested the effect of β-I on the response of Arabidopsis to this pathogen. For this purpose, we pre-treated Arabidopsis plants with β-I, followed by inoculation with *B.c.* Application of low (10 µM) or mid (50 µM) β-I concentrations did not negatively impact treated leaves on plants, while relatively high concentration of 1 mM β-I was toxic and led to necrosis (Fig. S10A). We also evaluated the effect of β-I (50 µM) on detached leaves of Arabidopsis plants infected with *B.c.* (Fig. 2A). We pre-treated the Arabidopsis plants twice (eight hours apart) with 50 µM β-I for 24 hours, which was followed by drop (5 µL) or spray inoculation with *B.c.* (2.5 x 10^5^ spores ml^-1^). Leaves treated with β-I were indistinguishable from the mock, however, they showed substantially reduced infection symptoms upon infection with *B.c.* (β-I+*B.c.*; Fig. 2A-B), compared to non-treated/non-infected controls (β-I or *B.c.*; Fig. 2A-B). This was evident from the measurement of the lesion size of infected leaves (Fig. 2C) and by qPCR quantification of *B.c.* infection, in which we determined by the quantification of fungal *Actin in planta* (Fig. 2D). We obtained similar results when we spray-inoculated Arabidopsis plants with *B.c.* (Fig. 2E). Taken together, these results indicate that pre-treatment of β-I can alleviate the effect of *B.c.* infection, leading to substantially reduced symptoms; thus, indicating a positive impact on plant response against this pathogen. To rule out that the observed reduction in infection is not caused by a direct inhibition of *B.c.* growth and general antifungal activity of β-I, we assessed fungal growth on agar plates supplemented with different concentrations of β-I. Our results demonstrated that *B.c.* growth was not affected at micromolar concentrations of β-I, but reduced the fungal growth at millimolar levels (Fig. S11), suggesting a plant-immunity triggering mechanism for increased resistance.

**Fig. 2.**
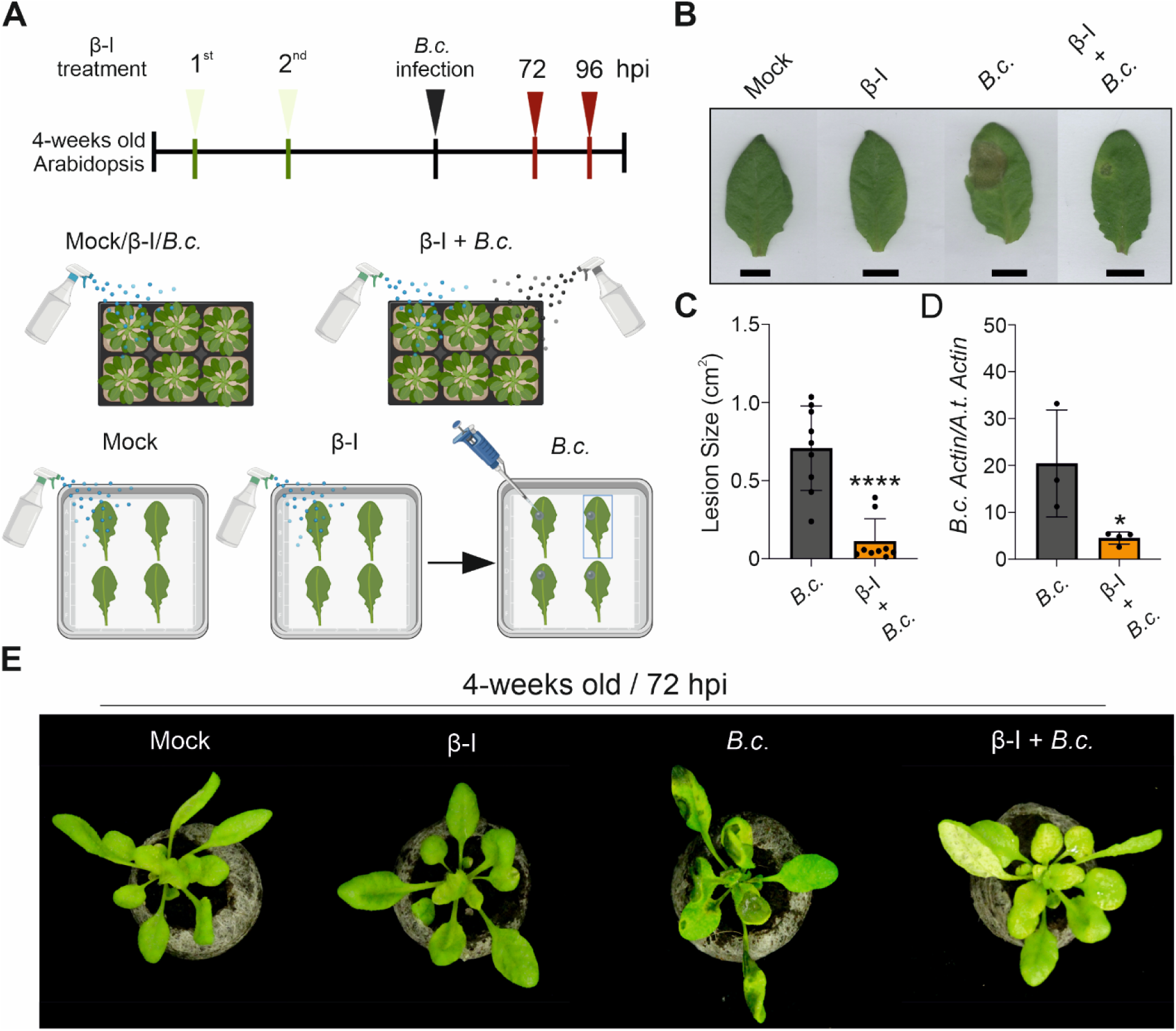
Application of β-ionone enhances resistance of Arabidopsis plants to *B. cinerea* infection. (*A*) Schematic representation of the experimental design for β-ionone (β-I) application and *B.cinerea* (*B.c.*) infection in detached leaves and intact plants. (*B*) Detached leaf assay of Arabidopsis wild type (Col-0) plants were treated with 1 % acetone (mock), 50 μM β-I, 5 μl drop inoculation with *B.c.*, or pre-treated with 50 μM β-I followed by 5 μl of *B.c.* drop inoculation (β-I + *B. cinerea*). β-I treatment was performed during the first 24 hours (with eight hours apart) prior to 5 μl drop inoculation with *B.c.* (2.5×10^5^ spores ml^-1^). (*C*) Lesion size quantification of detached leaves at 4 hours post inoculation (hpi) with 5 μl of *B.c.*. (*D*) qPCR quantification of *B. cinerea ActinA* gene relative to Arabidopsis *Actin* gene after β-I treatment and *B.c.* spray inoculation. (*E*) Photograph of Arabidopsis plants treated with 1 % acetone (mock), 50 μM β-ionone, spray inoculation with *B.c.* (2.5×10^5^ spores ml^-1^), and β-I + *B. cinerea*. The whole plant samples were pre-treated twice with 50 μM β-ionone followed by *B.c.* spray inoculation. Images were taken at 4 dpi. In (*B-C*), 10 rosette leaves were collected from 10 plants that were used for each experimental condition (*n* = 10). In (*D*), four biological replicates were used (*n*=4), and each sample was a pool of three leaves. Data represent single measurements, while bars and error bars represent the mean and ± SD. These experiments were repeated at least three times. Significance was calculated via an unpaired two-tailed Student’s *t*-test (\**p* < 0.05; \*\*\*\**p* < 0.0001). Figure (*A*) was prepared using Biorender.

### β-ionone treatment caused metabolic changes that enhances plant resistance to *Botrytis cinerea*

To determine the role of ABA, JA and SA in pathogen infection, we measured the content of these phytohormones in a time-resolved manner (Fig. 3A-C), following the same experimental design for β-I application and *B.c.* infection with minor modifications (Fig. S12). We also included a time point 0 which in fact represents ∼10-15 minutes after treatments due to the high amount of samples and treatments. This time point 0 was included to better dissect the pre-treatment with β-I in both β-I and β-I+*B.c.*, considering that β-I was sprayed 24 h before the time point 0 (see materials and methods and Fig. S12). Treatment with β-I reduced ABA content (but not other hormones) in Arabidopsis plants at early time points, most likely delaying the spreading of the infection. Similarly, ABA content was reduced at early time points in treated-plants with *B.c*. However, at the late time point (72 hours post infection (hpi)), when the infection is spreading throughout the whole plant and high amounts of ABA are needed, its content was ∼5-fold higher than in plants treated with the mock (Fig. 3A). By contrast, ABA content in β-I+*B.c.* treated-plants was ∼10-fold lower than in the *B.c.* treated-plants, suggesting a negative role of ABA in plant resistance to *B.c.*. Another key hormone, JA, was also increased at 72 hpi in *B.c.* infected plants, and, to a lower extent, in β-I+*B.c.* treated-plants, confirming the delayed infectious process in the β-I pre-treated plants (Fig. 3B). Interestingly, SA content was enhanced at 24 and 48 hpi in plants treated with either *B.c.* or β-I+*B.c.*, showing the highest increase (∼2-fold) at 72 h only in plants treated with β-I+*B.c.* (Fig. 3C). Additional evidence for the delayed infection was the production of camalexin which was 5- to 6-fold higher in *B.c.* treated-plants than in those treated with β-I+*B.c.* (Fig. S13) at 48 hpi. However, this difference is less pronounced (∼1.75-fold) at 72 hpi.

**Fig. 3.**
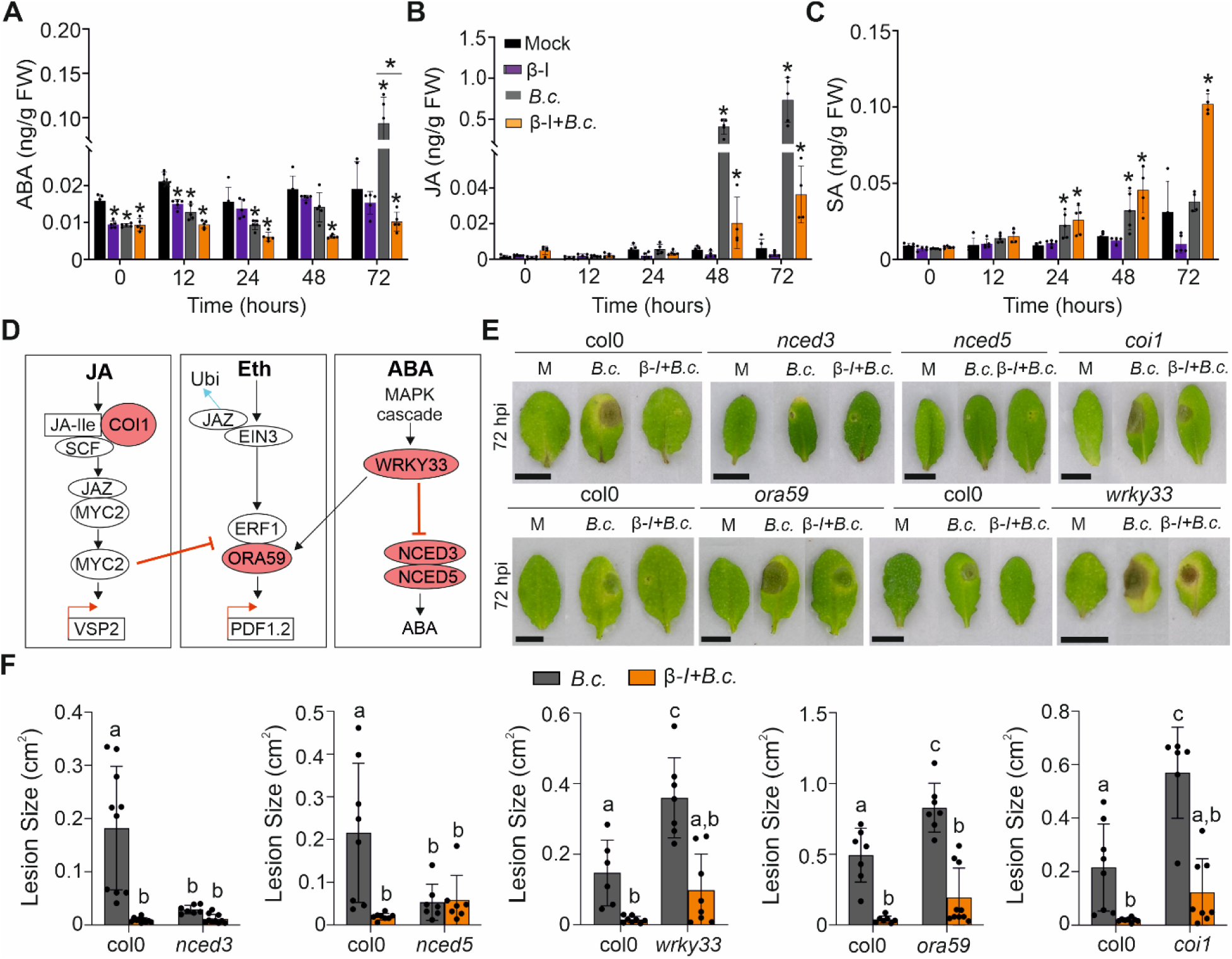
Involvement in hormonal signaling and general defense responses against *Botrytis cinerea*. (*A-C*) Time-resolved metabolic analysis of hormone (*A*) abscisic acid (ABA), (*B*) jasmonic acid (JA), and (*C*) salicylic acid (SA) contents in plants treated with β-ionone (β-I) and/or infected with *B. cinerea* (*B.c.*). Samples were collected at (0, 12, 24, 48, 72 h). Data points represent single measurements, while bars and error bars represent the mean and ± SD (*n=*5) for each treatment in each time point. Significance was calculated via Student’s unpaired, two-tailed t-test (\**p* < 0.05). Hormone quantification was performed once. (*D*) Simplified schemes depicting plant defense pathways against *Botrytis cinerea*. The scheme was prepared using previously published data (44-46, 80). (*E*) Detached leaf assay on 4-week-old plants of Arabidopsis from wild type (Col-0) and mutants altered in JA (*coi1*), Eth (*ora59*), ABA (*nced3* and *nced5*) and Eth-ABA-related (*wrky33*) pathways were treated with 1 % acetone mock (M), β-ionone (β-I), *B. cinerea* (*B.c.*), or β-ionone (β-I) prior to *B.c.* infection (β-I+*B.c.*). The 5 drop-inoculation of *B.c.* (2.5×10^5^ spores ml^-1^) method was used to infect plants while 50 μM β-ionone was sprayed. Col0, *nced3*, *nced5*, and *coi1* mutant leaves were treated and infected in the same plate, while *ora59* and *wrky33* were treated in separated plates, each of them with the respective wild type. (*F*) Lesion size of *B.c.* infection in detached leaves of Col-0 wild type and mutant plants. Data points represent single measurements, while bars and error bars represent the mean and ± SD (*n*=6-10). These experiments were repeated twice. Significance was calculated via ANOVA test with multiple comparisons (different letters represent significance, *p* < 0.05). Quantification of lesion size was performed using ImageJ software. Scale bar: 1 cm.

Plant defense against necrotrophs is orchestrated by crosstalk among plant hormones, such as JA, SA, ET, ABA, and brassinosteroids (BRs), which play a central role in plant defense against *B.c.* (Thomma et al., 1998; Audenaert et al., 2002; Lorenzo et al., 2003; Belkhadir et al., 2012; Denance et al., 2013; Kazan and Manners, 2013; He et al., 2017). To get a deeper insight into the role of JA, ET, and ABA (Fig. 3D), and to dissect the genetic components involved in the increased tolerance upon β-I application, we used the Arabidopsis mutant lines *coi1* (JA pathway), *ora59* (ET pathway) and *wrky33* (ET-ABA related) (Zheng et al., 2006; Sham et al., 2017), which were reported to show increased susceptibility to *B.c.* infection. In addition, *nced3* and *nced5* (ABA biosynthesis) mutants, which showed increased resistance to *B.c.* (Liu et al., 2015), were tested and compared to the wild-type. As expected, the latter mutants, containing lower ABA content compared with the wild type plants, showed increased resistance to the tested necrotrophic pathogen. However, the level of resistance did not increase upon β-I treatment (Fig. 3E-F). Application of β-I to the *coi1*, *ora59*, and *wrky33* mutants clearly increased their resistance to *B.c.*, as demonstrated by the significantly smaller size of the lesions (Fig. 3E-F).

### Transcriptome analysis of Arabidopsis plants upon β-ionone treatment and/or *Botrytis cinerea* infection

Based on our RNAseq analysis, phenotyping and hormone quantification, β-I enhances the resistance to *B.c.* most likely by affecting different defense pathways. Therefore, we performed RNAseq analysis on Arabidopsis plants pre-treated with β-I followed by *B.c.* infection (Datasets S11-S13). We compared the DEGs in Arabidopsis plants treated with β-I or infected with *B.c.* to determine the overlapping or specificities of gene expression. Interestingly, 36% (905) of the upregulated genes at 24 hpi with *B.c.* were also induced upon 24 hpt with β-I (Datasets S11-S12). These DEGs included defense response to fungus, immune response, immune system process and response to ABA stimulus (Fig. 4A; see the extended and full list in Fig. S14 and Dataset S14). Moreover, 44% (1138) of the downregulated genes after 24h of *B.c.* infection were repressed upon β-I treatment. Thus, there downregulated genes were related to biological processes, such as developmental process and photosynthesis (Fig. 4A; see the extended and full list in Fig. S15 and Dataset S15). These results suggest that β-I provokes a transcriptional response that overlaps with that triggered by *B.c.*, by reprogramming the expression of ∼2000 common DEGs. Thus, a pre-treatment with this compound would prepare the plant to respond in a better manner to the pathogen infection (Fig. 2). We also used MapMan software to depict these differences at the transcriptome level (Figs. S16-S17). Then, we characterized the kinetics of the changes at the transcript level for those plants that were pre-treated with β-I and then infected with *B.c.* at 24 and 48 hpi. In this case, 1412 and 1828 DEGs were commonly up- and down-regulated in Arabidopsis plants upon the treatment of β-I+*B.c.* at 24 and 48 hpi, respectively (Fig. 4A). The upregulated DEGs were grouped in GO biological processes such as defense response by callose deposition and cell wall thickening, defense response to fungus, and response to ABA/JA/SA (Fig. S18). By contrast, downregulated genes comprised GO biological processes such as cell growth, cell morphogenesis, development and photosynthesis (Fig. 4A; Fig. S19). In addition, we depicted changes in metabolic processes (metabolism overview) and stress response (biotic stress) using the software MapMan (Figs. S20-S21). We noticed that the upregulated genes were predicted to localize in the nucleus (24%), cytosol (19%), plasma membrane (17%) and chloroplast (11%); while for the downregulated ones were in the nucleus (24%), chloroplast (17%), plasma membrane (16%) and extracellular space (13%; Fig. 4B and Datasets S16-S17). Next, we analyzed the expression pattern of 49 key genes related to plant defense/response to *B.c.*. These genes are involved in signaling, cell wall, hormone metabolism, biotic stress, and encode transcription factors (Fig. 4C). Interestingly, β-I, *B.c.* and β-I+*B.c.* treatments caused the relatively similar effect on the expression of 29 of these genes (26 induced and 3 repressed) (Fig. 4C). The remaining 20 genes showed an opposite response to β-I treatment, compared to *B.c.* or β-I+*B.c.* infection (Fig. 4C). Thus, genes that respond similarly to all treatments *i.e.,* β-I, *B.c.* and β-I+*B.c.*, are possibly involved in plant’s defense response against *B.c.*, suggesting that β-I mimics *B.c.* infection.

**Fig. 4.**
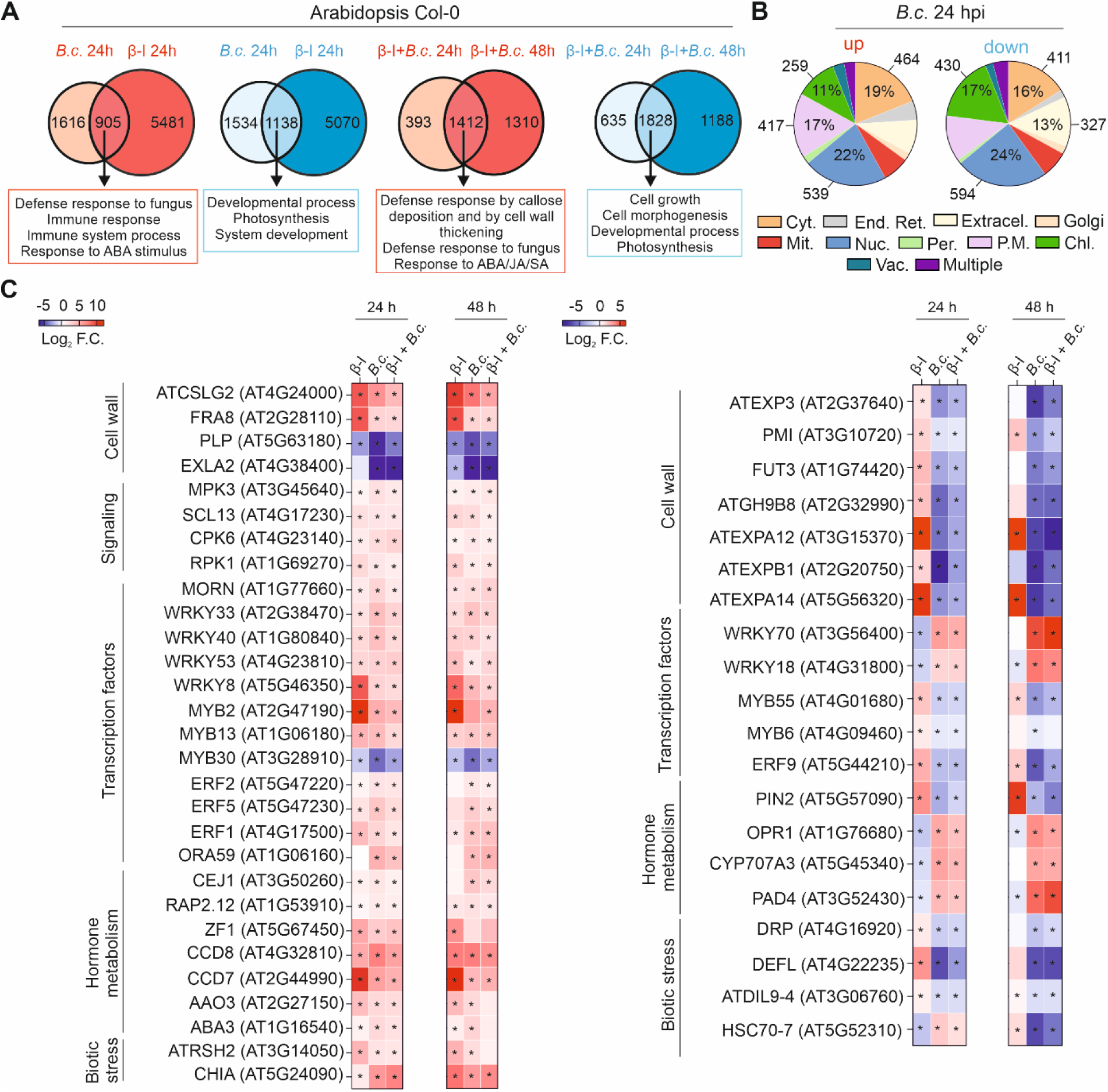
RNA sequencing (RNAseq) analysis of plants pre-treated with β-ionone and infected with *B. cinerea*. (*A*) Venn diagrams of DEGs identified through RNAseq in Arabidopsis plants infected with *B. cinerea* (*B.c.*) and pretreated with β-ionone followed by *B. cinerea* infection (β-I+*B.c.*) at 24 and 48 hpi. Venn diagrams representing up- and downregulated genes are shown in red and blue, respectively. Some of the most important GO biological processes enriched in the overlapping genes are shown (for full enriched GO list see Table S1 and Fig. S4). (*B*) Distribution of cellular compartments for differentially expressed genes (DEGs, up and down) present after 24 h of *B.c.* treatment. Subcellular localization analysis was performed in SUBA4 online software (https://suba.live/). Genes encoding proteins with 2 or more localizations were grouped in the “multiple” category. Each color in the pie chart represents a cellular compartment. (C) Heatmap representation of changes at transcript level of Arabidopsis plants treated with β-I, *B.c.*, and β-I + *B.c.* at 24 and 48 hpi. Heatmaps show the common up- and downregulated genes involved in biological processes such as cell wall biosynthesis and defense mechanisms, signaling pathways, transcription factors, hormone metabolism, and biotic stress. Statistical analysis for the RNAseq was performed using DESeq2 with Benjamini and Hochberg’s approach for controlling False Discovery Rate (FDR). Genes were adjusted Log2 fold change expression (*padj*<0.05). Cyt: cytoplasm; End. Ret.: Endoplasmic reticulum; Extracel.: extracellular; Mit.: mitochondria; Nuc.: nucleus; Per.: peroxisome; P.M.: plasma membrane; Chl.: chloroplast; Vac.: vacuole.

Genes with the opposite response to β-I treatment, compared with *B.c.* and β-I+*B.c.* infection may be needed for proper *B.c.* infection. Genes with the similar expression pattern are involved in cell wall synthesis (e.g., *CELLULOSE SYNTHASE LIKE G2* (*CSLG2*)), signaling pathways (e.g., *MPK3*), hormone metabolism (e.g.*, CAROTENOID CLEAVAGE DIOXYGENASES 7* (*CCD7*) and *CCD8*), and biotic stress (*CHITINASE A* (*CHIA*)), or encode transcription factors (e.g., *WRKY*s and *ERF*s). β-I treatment upregulated Arabidopsis genes that are associated with cell wall defense response and loosening (e.g., *EXP3*), signaling pathways (e.g., *CALCIUM-DEPENDENT PROTEIN KINASE* 31 (*CPK31*)), transcription factors encoding genes (e.g., *MYB*s), hormone metabolism (*PIN-FORMED 2* (*PIN2*)), and biotic stress-related genes (e.g., *DEFENSIN-LIKE PROTEIN* (*DEFL*)), while they were downregulated in response to *B.c.* and β-I+*B.c.-*treated plants (Fig. 4C). Moreover, β-I treatment caused downregulation of genes coding for transcription factors (e.g., *WRKY70* and *WRKY18*), or that were involved in hormone metabolism (*2-OXOPHYTODIENOATE REDUCTASE 1* (*OPR1*) and *CYP707A3*), and biotic stress response (*PHYTOALEXIN DEFICIENT 4* (*PAD4*) and *HEAT SHOCK PROTEIN 70-7* (*HSC70-7*)), while they were upregulated in response to *B.c.* and β-I+*B.c.* treatment (Fig. 4C).

### β-ionone effect is conserved in crop plants

We evaluated whether β-I can also increase resistance to *B.c.* in other dicotyledonous crops, considering the large yield losses and economic impact of this pathogen. Therefore, we evaluated the protective role of β-I in several cultivars of the cash crop, tobacco (*Nicotiana tabacum*), and the edible crop, tomato (*Solanum lycopersicum*). We followed an experimental setup designed for β-I treatment and *B.c.* infection as described in Fig. S3. First, we tested the effect on detached leaves of two tomato cultivars (IPA6+ and MaxiFort). Our results showed larger lesions in the *B.c.* infected leaves than in β-I+*B.c.*-treated leaves in both cultivars (Fig. 5A, B), indicating less severity in tissue maceration by *B.c.* upon β-I treatment. In tomato, *B.c.* infection in β-I+*B.c.*-treated leaves was lower than in leaves infested with *B.c.* without β-I treatment (Fig. 5C). In order to rule out any biological response that could interfere with our assay after cutting the tomato leaves, we also treated intact tomato and tobacco plants and observed similar response (Fig. S3). On one hand, control plants that were sprayed with mock exhibited a normal green phenotype, while the *B.c.* infected plants showed a severe gray mold infection including necrotic lesions and tissue maceration. On the other hand, plants treated with β-I+*B.c.* looked healthy and showed only few and small necrotic lesions on leaves (Fig. 5D). This phenotypic difference mirrored ∼100-fold lower presence of *B.c.* in β-I+*B.c.* plants compared to *B.c.* treated-plants (Fig. 5D). Similarly, tomato plants inoculated with *B.c.* looked weaker than control plants, showing bent branches with a sever necrotic lesions. By contrast, β-I+*B.*c-treated plants looked more vigorous and developed only few necrotic lesions on their leaves (Fig. 5E). We also evaluated the effect of β-I on tomato fruits using the Micro-Tom variety due to its smaller size and much shorter life cycle than the previously used cultivars (Fig. S3). After 7 days post harvesting (dph), non-treated tomato fruits were normally red colored and had a smooth skin, while fruits infected with *B.c.* showed fungal growth in the sepals and in the skin at 7 dpi (Fig. 5F). Surprisingly, tomato fruits pre-treated with β-I did not develop disease symptoms and showed extremely reduced levels of fungal growth (Fig. 5F). To confirm the effect of endogenous β-I on *B.c.* infection, we used fruits of two transgenic tomato lines of the varieties Red Setter/R.S. and IPA6+, which express the *LYCOPENE β-CYCLASE* (*LCYB*) gene from tomato (H.C.) or daffodil (pNLYC#2) and contain 60% and 100% higher β-I content, compared to their respective wild type (Mi et al., 2022). Tomato wild type R.S. and IPA6+ plants showed red-colored fruits and smooth and firm skin after 7 dpi. Infection with *B.c.* led to damaged skin with necrotic lesions (Fig. 5G). Fruits of *LCYB*-expressing tomato plants were orange-colored with smooth skin (upper panel). Upon *B.c.* infection, these fruits showed small necrotic lesions in H.C. and pNLyc#2, but maintained their firmness (Fig. 5G). This indicates a biological role of endogenous β-I in alleviating the effect of *B.c.* infection in plants.

**Fig. 5.**
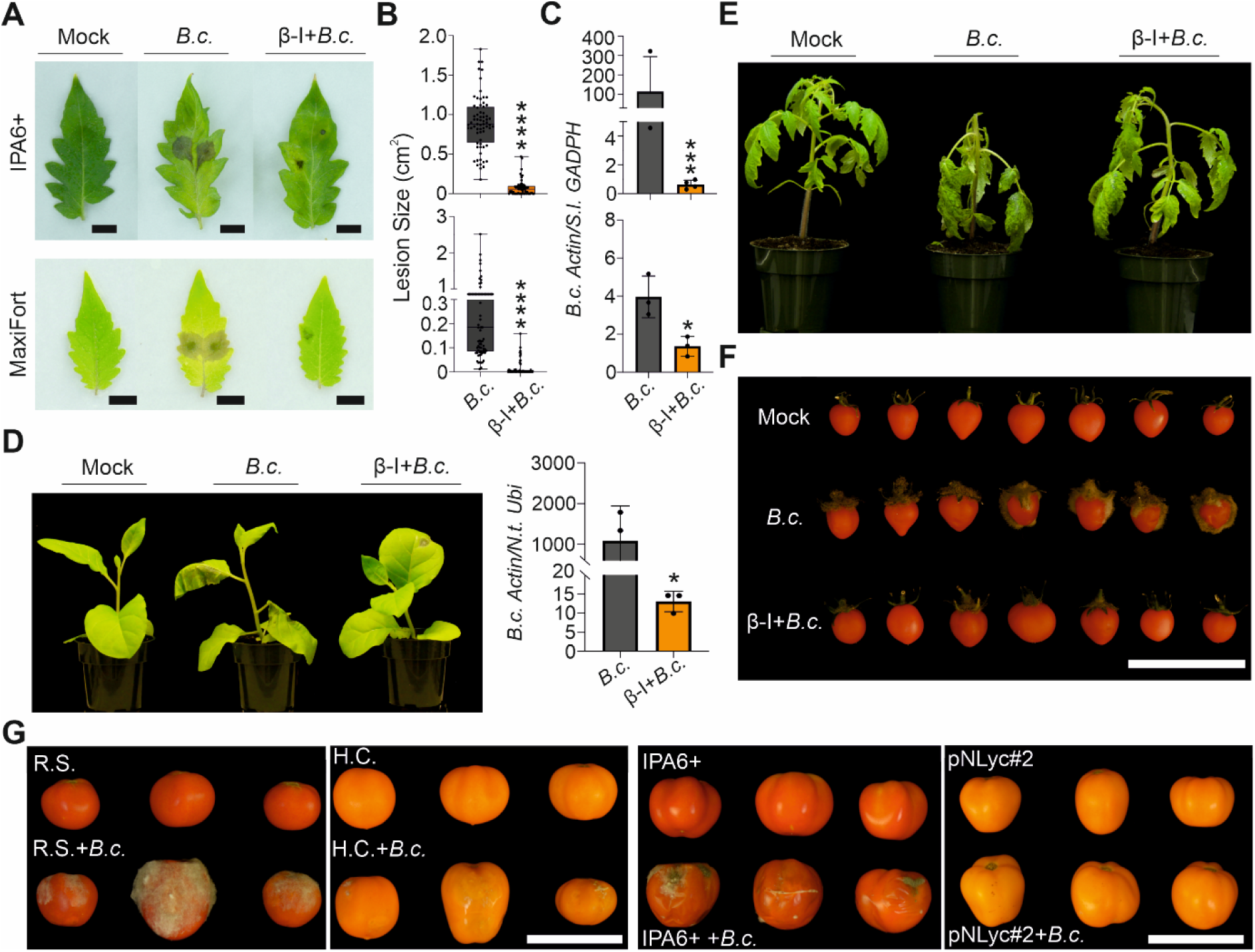
Effect of β-ionone on tomato and tobacco plants. (*A*) detached leaves of tomato plants (cv. IPA6+ and Maxifort) that were treated with 1% acetone (mock), 5 μl drop inoculation of *B. cinerea* (*B.c.*) using 2.5×10^5^ spores ml^-1^, and 50 μM β-ionone (β-I) application twice during 24 h (eight hours apart) followed by 5 μl drop inoculation *B. cinerea* (β-I+*B.c.*). (*B*) Lesion size and (*C*) fungal content in detached leaves of tomato (cv. IPA6+ and Maxifort) in β-I and/or *B.c.* treatments. Amplification of *B. cinerea ActinA* relative to tomato housekeeping gene *GADPH* was quantified in leaves treated with *B.c.* or β-I+*B.c.* at 72 hours post inoculation (hpi). (*D*) Phenotyping and *B.c.* quantification in *Nicotiana tabacum* (cv. Xanthi) in *B.c.* and in β-I+*B.c.* treated plants (72 hpi). Transcript levels of *B.c. ActinA* relative to the *Nicotiana tabacum Ubiquitin* (*N.t. Ubi*) gene were quantified in *B.c.* and β-I+*B.c.* treated plants. Significance in *B-D* was calculated using an unpaired two-tailed Student’s *t*-test (\**p* < 0.05, \*\*\**p* < 0.001, **** *p* < 0.0001). (*E*) Disease symptoms in tomato plants (cv. IPA6+) treated with *B.c.*, or β-I+*B.c.*. Photos were taken at 72 hpi. (*F*) Tomato fruits (cv Micro-Tom) treated with 1% acetone (Mock), 5 μl of drop inoculation with *B.c.*, or 50 μM β-I application twice during 24 h (eight hours apart) followed by 5 μl of drop inoculation with *B.c.* (β-I+*B.c.*). Red fruits and green sepals were observed in the mock treatment; while fungal growth was observed on fruits inoculated with *B.c.*. In β-I+*B.c.* treatment, fungal growth was limited on tomato fruits at 7 days post inoculation (dpi). (*G*) Tomato fruits of wild type (IPA6+ or Red Setter) and transgenic plants overexpressing lycopene β-cyclase with and without spray inoculation with *B.c.*. Fruits that were not inoculated with *B.c.* remained healthy and showing no symptoms of gray mold disease at 7 dpi. Tomato wild type plants cv. Red Setter (RS) and IPA6+ inoculated with *B.c.* showed fungal hyphal growth on their fruits. On the other hand, the orange-colored fruits in plants expressing *LCYB* gene that produced 60 and 100% higher β-I content in H.C. and pNLyc#2 transgenic plants, respectively, than in their corresponding wild type plants showed small necrotic lesions on their skin. Scale bar: 10 cm.

## Discussion

*B.c.* is a necrotrophic fungus that causes gray mold disease on a wide range of plant species. It is responsible for pre- and post-harvest decay of fruits and vegetables in greenhouses, open fields, and during storage (Dean et al., 2012; AbuQamar et al., 2017). *B.c.* infests economically important crops, such as tomato, and ornamental flowers, causing losses in the range of 15-40% due to postharvest spoilage (Legard et al., 2000). So far, the only mean to manage gray mold disease is the application of synthetic fungicides; however, *B.c.* has developed resistance to these chemicals. In addition, chemical fungicides may affect human health and have negative impact on the environment. Here, we identified the scent apocarotenoid β-I as a signaling molecule that provokes Arabidopsis biotic stress response and enhances plant defense against *B.c.*. But, how does β-I trigger these effects at molecular and phenotypic level?

β-I is naturally produced in plastids through the non-enzymatic oxidation of β-carotene or CCD-catalyzed cleavage. Interestingly, β-I shares many features with another plastid volatile signaling molecule, the apocarotenoid β-cc, which triggers the expression of hundreds of nuclear-encoded genes (Ramel et al., 2012; D’Alessandro et al., 2018); thus, resulting in enhanced plant tolerance to high light-induced oxidative stress (Ramel et al., 2012) and abiotic stresses (D’Alessandro et al., 2019). The two compounds differ in their chain-lengths and the nature of the carbonyl group, but both of them i) are volatiles synthesized in plastids, ii) share the same precursor (β-carotene), iii) can be produced enzymatically or non-enzymatically induced by high light. Therefore, we hypothesized that β-I might be also a bioactive apocarotenoid. Our results showed that β-I triggered the expression of thousands of nuclear-genes, suggesting that β-I is a signaling molecule. Plastids rely on signals from the nucleus to coordinate their gene expression and adjust their biochemical and other biological processes to the status of the cell. Depending on their needs, plastids, however, also generate retrograde signals that regulate nuclear gene expression (Nott et al., 2006; Woodson and Chory, 2008; Chan et al., 2016; de Souza et al., 2017). Because β-I is produced in plastids and modulates the expression of nuclear genes, it can be considered as a novel retrograde signal. Here, we showed that β-I-induced genes are involved in plant defense to pathogens, and repressed genes involved in the biosynthesis of tetrapyrroles, chlorophylls, and isoprenoids, and in photosynthesis (Fig. 1A, C; Fig. S9; Dataset S2). The reduction in the biosynthesis of photosynthetic pigments and the perturbation of plastid gene expression initiate a retrograde control of the expression of photosynthesis-associated nuclear genes (*PhANGs*) (Barajas-Lopez et al., 2013; Chan et al., 2016; Hernandez-Verdeja and Strand, 2018; Wu and Bock, 2021). Prominent *PhANGs* responsive to retrograde signals are genes encoding LHCBs and RBCS. In addition, *GUN* genes are involved in retrograde signaling control of gene expression in the nucleus (Wu and Bock, 2021). In line with these findings, we observed a massive reduction in the transcript levels of all *LHCBs* and *RBCS*, as well as in *GUN4* and *GUN5* genes upon the application of β-I (Fig. 1C; Fig. S9B; Dataset S2), thus, supporting the retrograde signaling function of β-I. Recently, Mitra *et al*. (54) reported on the involvement of β-cc in plant defense against herbivores. Upon herbivory attack or exogenous treatments, β-cc binds to the key MEP pathway enzyme Deoxyxylulose 5-phosphate synthase (DXS), reducing its activity and, hence, the MEP pathway flux and the biosynthesis of isopentenyl diphosphate (IPP) and dimethylallyl diphosphate (DMAPP) and thereof derived isoprenoids. Indeed, this decrease was reflected in lower chlorophyll and carotenoid contents. In addition, β-cc increment enhanced the content of the 2-C-methyl-D-erythritol-2, 4-cyclodiphosphate intermediary in the cytosol, which was reported to be a signaling molecule that upregulates SA signaling (Lemos et al., 2016; Onkokesung et al., 2019) and enhances plant defense (56). In the current study, exogenous application of β-I reprogrammed the transcriptome from growth to defense mode. However, it remains unclear if *B.c.* infection causes an increase in the formation of β-I. Nevertheless, the high ROS levels arising in cells upon *B.c.* penetration and infection (Heller and Tudzynski, 2011; Torres et al., 2013; Rossi et al., 2017) could break β-carotene into β-I, arguing in favor of the production of this compound upon infection.

Arabidopsis defense mechanisms against *B.c.* occur at several cellular levels in distinct cellular compartments including the cell wall, plasma membrane, cytoplasm, and the nucleus (Fig. 6) (AbuQamar et al., 2017). We found 24 genes in our transcriptome, which overexpression or downregulation were previously reported to confer full immunity against *B.c.* (Fig. 6; Table S2). These genes encode different types of proteins, including lipid transfer proteins (LTP), peroxidases (PER), proteinase inhibitors (PIs), plasma membrane receptors, polygalacturonase inhibiting proteins (PGIPs), and map kinases (MAPKs; Table S2). The encoded proteins are involved in cuticle permeability (e.g., BODYGUARD/BDG) (Sieber et al., 2000; Kurdyukov et al., 2006; Chassot et al., 2007; Serrano et al., 2014), induced systemic resistance (TLPs) (Arondel et al., 2000), response to (oxidative) stress (PERs) (Tognolli et al., 2002), response to fungus and wounding (PIs) (Dunaevskii Ia et al., 2005), plasma membrane receptors (BAK1 and SOBIR1) (Zhang et al., 2013), inhibition of polygalacturonases (PGs) (De Lorenzo et al., 2011), and signal transduction (MAPKs) (Ren et al., 2008; Pieterse et al., 2009; Fiil and Petersen, 2011; Galletti et al., 2011; AbuQamar et al., 2017) (Table S2). Surprisingly, the increase/decrease in expression of all these genes in our transcriptome was in line with the previously reported Arabidopsis enhanced resistance to *B.c.* in the respective mutants (Table S2), suggesting their contribution to the observed tolerance against *B.c.* upon β-I application (Fig. 2B, E). Interestingly, we also observed very high expression of other members of PER (PER32, 42-fold; PER50, 6.4-fold), PI (*KTI/At1g73260*, 1303-fold), and LTP (*At4g12520*, 485-fold; *AZI5*/*At4g12510*, 1006-fold; *At1g18280*, 168.4-fold; *At4g12500*, 196-fold; LTP2, 284-fold) families (Fig. 6; Table S3), indicating that the overexpression of these genes might also contribute to the β-I-induced resistance against *B.c*.

**Fig. 6.**
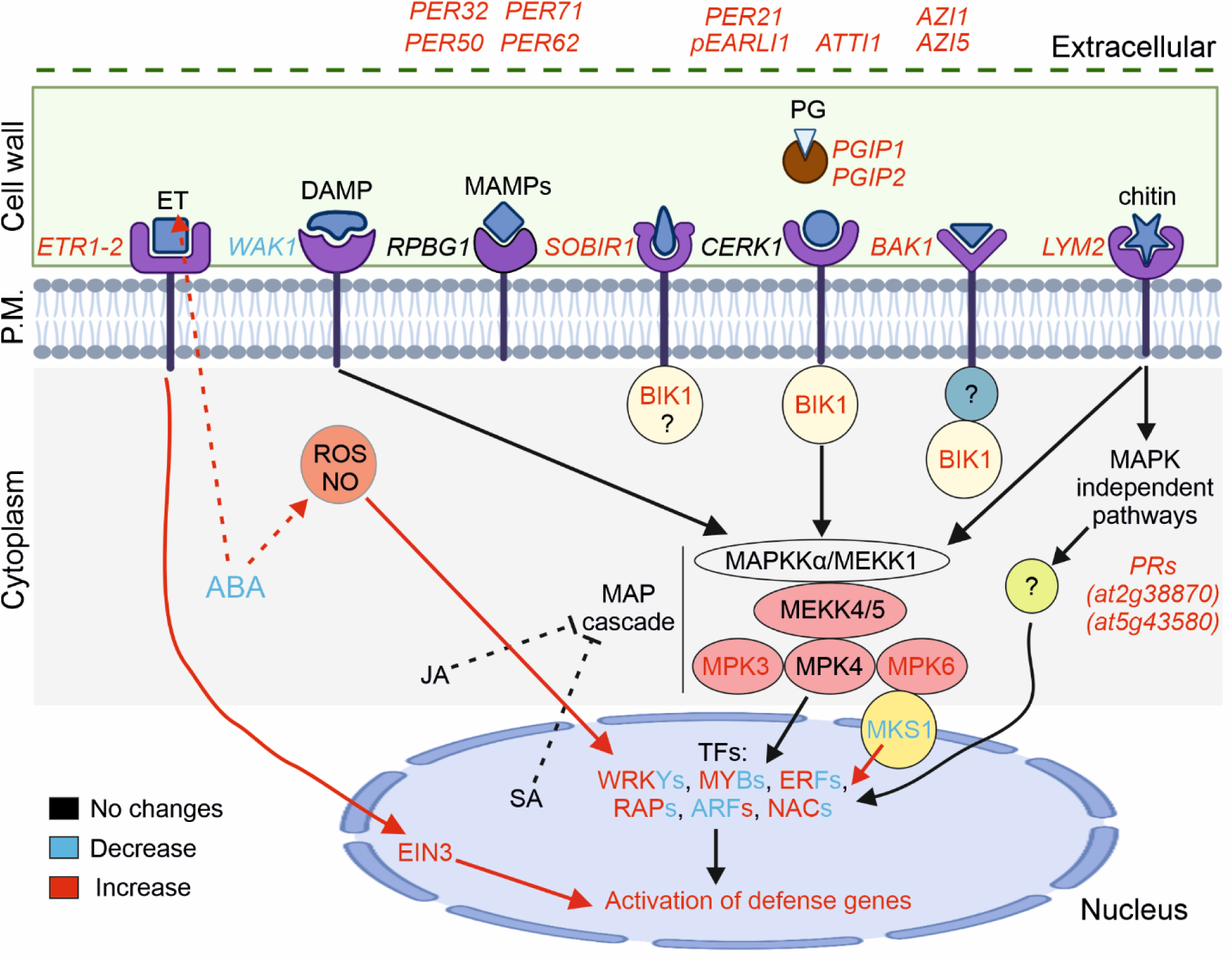
Schematic representation of β-ionone effect at molecular and metabolic level in Arabidopsis. Schematic model depicting molecular processes required to activate plant defense against *B.c*. Genes that were up- and down-regulated upon β-I treatment (24hpt) which contribute to enhanced *B.c.* resistance are shown in red and blue, respectively. Arabidopsis transgenic lines with higher/lower transcript level of these genes were previously reported to positively contribute with enhanced resistance against *B. cinerea*. Question marks represent unknown proteins or steps upon *Botrytis* infection. P.M.: plasma membrane; PRs: pathogenesis related proteins; MAPK: mitogen activated protein kinase; TFs: transcription factors (adapted from AbuQamar *et al*., 2017). The figure was prepared using Biorender.

Pathogen attack stimulates the synthesis of phytohormones such as SA, JA and ET that regulate specific immune responses (Glazebrook, 2005; Pieterse et al., 2009). Both SA and JA/ET pathways are involved in response to biotrophic and necrotrophic pathogens, respectively (Antico et al., 2012; Toth et al., 2016). Other phytohormones, such as ABA and brassinosteroids, regulate plant immunity, mainly by interacting with transcription factors, or through camalexin biosynthesis and callose deposition (Audenaert et al., 2002; Denance et al., 2013). ABA has a negative effect on defense against *B.c.* (Audenaert et al., 2002; Windram et al., 2012), and it is needed for fungal colonization and for spreading the infection across the plant (Audenaert et al., 2002; Schmidt et al., 2008). We observed a decrease (up to 24 hpi) and an increase (72 hpi) in ABA content at early and late time points, respectively, in response to *B.c*. These changes are in line with previous data showing the highest ABA accumulation at later time points (more than 40 hpi) and a slight reduction at early time points (12-18 h) in Arabidopsis (Liu et al., 2015; Liu et al., 2017). Our results showed a decrease in ABA content upon β-I treatment at early time points and a constantly reduced ABA content across all time points in the β-I+*B.c.* treatment (Fig. 3A). Interestingly, the ABA content remained ∼10-fold lower in β-I+*B.c.*-treated plants than in those infected with *B.c.* at 72 hpi; thus, enhancing resistance against *B.c*. In previous studies using Arabidopsis and tomato ABA-deficient mutants, several mechanisms have been proposed for the higher resistance against *B.c.*, including induction of ROS and nitric oxide (NO) and increase permeability of the cuticle (Asselbergh et al., 2007; L’Haridon et al., 2011; Sivakumaran et al., 2016). In Arabidopsis, NO induces ET biosynthetic genes and ET production in response to *B.c.* (Mur et al., 2012). Therefore, ABA reduces plant resistance to *B.c.* probably through the reduction of NO levels and suppression of both ROS and ET production (AbuQamar et al., 2017). In the present study, reduced ABA might have caused enhanced NO, ROS and ET levels allowing the activation of defense genes (Fig. 6). In addition, ABA deficiency increases cuticular permeability and resistance to *B.c.* as observed in the tomato *sitiens* and the Arabidopsis *abi2* and *abi3* mutants (Curvers et al., 2010; L’Haridon et al., 2011). In line with these findings, we did not observe an increase in *B.c.* resistance upon β-I treatment in the ABA-deficient *nced3* and *nced5* mutants, while β-I treatment of *ora59* (ET), *coi1* (JA), and *wrky33* (ET/JA, ABA, and other responses) restored the susceptibility of these mutants to the wild-type level (Fig. 3D-F). This suggests that β-I might interfere with the ABA biosynthesis to lower ABA content. In addition, several studies highlighted the crosstalk of SA, JA, and ET with ABA in regulating plant defense against *B.c.* (De Bruyne et al., 2014; Jiang et al., 2016). Although we did not observe an increase in their content upon β-I application, enhanced JA and SA level at later time points in β-I+*B.c* treatment might also contribute to defense against *B.c*..

We also evaluated if the effect of β-I is conserved in other model plants. Infection experiments in wild type cultivars of tobacco and tomato revealed a similar positive effect on resistance against *B.c.*. Moreover, transgenic tomato fruits with enhanced β-I content exhibited increased resistance to *B.c.*. These results point to a fairly conserved mechanism in *Brassicaceae* and *Solanaceae*, both dicotyledonous species and main targets of *B.c.*. In addition, time-course infection experiments in two tobacco cultivars (*Xanthi* and *Petit havana*) revealed that β-I treatment might delay plant decay by 6-9 dpi (Fig. S22).

Taken together, our results showed that β-I is a signaling molecule that provokes a transcriptional response overlapping with that caused by *B.c.* and following a defense-growth trade-off. Moreover, we demonstrated that this apocarotenoid could enhance Arabidopsis resistance to *B.c*, likely via modulating hormonal contents and regulating the expression of genes involved in plant defense, which is also conserved in tobacco and tomato. These findings uncover a new member of the apocarotenoid family of hormones and regulatory metabolites and open up the possibility of developing a bio-fungicide that could replace the heavy use of chemical fungicides in field and post-harvesting.

## Materials and Methods

### Plant genotype and growth conditions

*Arabidopsis thaliana* Col-0 seeds and other mutant genotypes were grown on Jiffy soil (Jiffy Product International AS, Norway). They were placed at 4°C for three days before being transferred to a growth chamber with 16 h light/8 h dark and 180 umol/m s illumination at 21°C day/18°C night for four weeks. Tobacco (*Nicotiana tabacum* cv. *Xanthi NN* and *N. tabacum* cv. *Petit Havana*) and tomato (*Solanum Lycopersicum L.* cv. IPA6+) wild type plants were grown under greenhouse conditions with 16 h light/8 h dark and 180 umol/m s at 28°C. For *B. cinerea* assays, Arabidopsis, tobacco and tomato wild type seeds were sown in individual pots and grown for 4-6 weeks depending on the species. *B. cinerea* infection in detached tomato leaves was performed using 10-week-old F1 plants of the interspecific hybrid ‘Maxifort’ (*Solanum lycopersicum* L. × *Solanum habrochaites* S. De Ruiter, Bergshenhoek, The Netherlands). For fruit infection experiments, we used greenhouse-harvested tomato fruits from cultivars Micro-Tom and transgenic pNLyc#2 (cv. IPA6+) and H.C. lines (cv. Red Setter) (Apel and Bock, 2009; Mi et al., 2022). All tomato plants were grown under greenhouse conditions, with scheduled 20-20-20 fertilization once a week.

### *B. cinerea* inoculation of Arabidopsis plants

For pathogen assays, detached leaves from four-week-old Arabidopsis plants were used to perform infection experiments using *B. cinerea*. In addition, intact plant infection experiments were also performed to avoid any hormonal perturbation in plants. First, detached leaves were sprayed twice within 24 hours (at 16 and 24 h prior *B. cinerea* infection) with 1% acetone (mock) or β-ionone (50 μM; dissolved in 1% acetone). These plants were drop-inoculated with 5 µl *B. cinerea* conidial suspension containing 2.5×10^5^ spores ml^-1^ or spray-inoculated with *B. cinerea* (2.5×10^5^ spores ml^-1^) on the whole plants. Arabidopsis plants sprayed twice with β-ionone within 24 hours with 1% acetone or β-ionone (50 μM; dissolved in 1% acetone and 0.05% tween 20) were used as the control. To establish the disease for drop inoculation, detached leaves were placed together with moist Whatman filter paper on 245 mm square bioassay dishes and sealed with parafilm. Treated/infected plants were covered with a plastic dome and sealed with tape to maintain high humidity. All the experimental groups were kept under the same conditions for 3-4 days at room temperature under dark conditions. For Arabidopsis metabolites and LC-MS based hormone quantification, samples were collected at 0, 12, 24, 48, and 72 hours. Five biological replicates were used and each sample contained ∼250 mg of tissue (fresh weight).

### Lesion size measurements

Arabidopsis Col-0 seeds were sown in Jiffy soil and grown in a growth chamber under controlled climate conditions as described above. Mature rosette leaves were detached from four-week-old plants and subjected to treatments or infection as described above. Three to four days post-inoculation (dpi), lesion size was measured. Photographs of the infected plants with the lesions were taken and lesion size measurements were quantified using ImageJ 1. x software (Schneider et al., 2012).

### RNA extraction

Total RNA was prepared from Arabidopsis, tobacco and tomato plant material. Plant samples were collected from leaf tissue and placed in 2 ml microcentrifuge tubes together with three steel beads (2.3-mm diameter), further frozen in liquid nitrogen and grounded for 30 seconds (Mini-Beadbeater-96, #1001, Biospec Products). For the RNAseq experiment, plant material was collected following the experimental design described in Fig. S2. Total RNA was extracted using the Direct-zol RNA Miniprep Plus Kit (Zymo Research according to the manufacturer’s instructions; see Methods S1).

### RNAseq analysis of Differentially Expressed Genes (DEGs)

The analysis of DEGs was performed between two conditions, β-ionone treated Arabidopsis plants at 3 h *vs*. 24 h, β-ionone at 24 h *vs. B. cinerea* at 24 h, and β-ionone + *B. cinerea* treated plants at 24 h *vs.* 48 h (three biological replicates per control) was performed using DESeq2 R package (Yu et al., 2012). DESeq2 provides statistical routines for determining differential expression in digital gene expression data using a model based on the negative binomial distribution (Dai et al., 2021). The resulting *p* values were adjusted using the Benjamini and Hochberg’s approach for controlling the False Discovery Rate (FDR). Genes with an adjusted *p* < 0.05 found by DESeq2 and at least an increase of 20% and a reduction of 35% were assigned as differentially expressed.

### Gene expression analysis by qPCR

cDNA was synthesized from 1 µg of the total RNA using iScript Kit (BIO-RAD Laboratories, Inc, 2000 Alfred Nobel Drive, Hercules, CA; USA) according to the manufacturer’s instructions. The *B. cinerea* DNA was quantified in infected plants by qPCR analysis, based on the relative expression of *B. cinerea ActinA* (*BcActinA*) to the Arabidopsis (*Actin*), tobacco (*Ubiquitin*) and tomato (*GADPH*) housekeeping genes (Table S1). Expression analysis was analyzed using gene-specific primers (Table S1). The statistical significance was determined via Student’s unpaired, two-tailed *t*-test using GraphPad Prism software. The data are shown as means and the error bars representing the ± standard deviation (SD) of four independent biological replicates. The mean values showing asterisks are significantly different from the corresponding control (*P*<0.05).

### Quantification of plant metabolites and hormones

For the quantification of endogenous metabolites and hormones, about 250 mg (fresh weight) of grounded Arabidopsis leaves, spiked with internal standards, i.e., 1 ng of D_3_-β-ionone, 1 ng of D_6_-ABA, and 10 ng of D_4_-SA, were extracted with 1.5 mL of methanol containing 0.1% BHT in an ultrasound bath for 15 minutes. After centrifugation at 13000 rpm and 4°C for 8 minutes. The supernatant was collected and kept at -20 °C. The residue was re-extracted with 1 mL of 10 % methanol with 1% acetic acid in an ultrasound bath for 5 minutes, followed by incubation on ice under shaking at 500 rpm for 45 minutes. After centrifugation at 4000 rpm at 4 °C for 8 minutes, the two supernatants were combined and filtered by using a 0.22 µm filter. UHPLC-MS/MS analysis of plant metabolites and hormones was performed on a Dionex Ultimate 3000 UHPLC system coupled with a Q-Orbitrap-MS (Q-Exactive plus MS, Thermo Scientific) with a heated-electrospray ionization source according to (Mi et al., 2019; Jia et al., 2021).

### *B. cinerea* infection of tobacco and tomato plants

For infection experiments in tobacco and tomato we followed the same protocol as for Arabidopsis but with slight modifications for leaves and fruits (see Methods S1).

### Statistical Analyses

The statistical analyses were performed in “R” (RNAseq) or with the software GraphPad Prism 9.0 employing *t*-test or ANOVA test depending on the experimental design of each experiment. The tests are described within Materials and Methods section for each experiment and also in the legend of each figure.

## Acknowledgements

We are grateful to Kit Xi Liew for his assistance in the quantification of hormones and metabolites, Dr. Justine Braguy, Dr. Jian You Wang for their advice and discussions, Lamis Berqdar for her technical support. Funding for this work was provided by Khalifa Center for Genetic Engineering (KCGEB) (31R286 to Sy. A.) and baseline BAS/1/1039-01-01 given to Sa. A. from King Abdullah University of Science and Technology (KAUST).

## Author Contributions

Sy. A. and Sa. A. conceived the project; A. F., J.C.M., Sh. A., A.S. performed the Arabidopsis inoculation studies. J.C.M conceived the tomato and tobacco study and performed it with A.F. J.M. quantified the hormones. A.F. and J.C.M. performed the expression analysis and the RNAseq experiments. A. F. and J.C.M. analyzed the data. J.C.M. and A. F. wrote the paper. Sh. A., Sy. A., and Sa. A. revised the paper.

## Data availability

All data is included in the main body or supplemental information.

